# Validation of chronic restraint stress model in rats for the study of depression using longitudinal multimodal MR imaging

**DOI:** 10.1101/2020.03.20.998195

**Authors:** Bhedita J Seewoo, Lauren A Hennessy, Kirk W Feindel, Sarah J Etherington, Paul E Croarkin, Jennifer Rodger

**Affiliations:** Experimental and Regenerative Neurosciences, School of Biological Sciences, The University of Western Australia, WA 6009, Australia; Brain Plasticity Group, Perron Institute for Neurological and Translational Science, WA 6009, Australia; Centre for Microscopy, Characterisation & Analysis, Research Infrastructure Centres, The University of Western Australia, WA 6009, Australia; School of Biomedical Sciences, The University of Western Australia, WA 6009, Australia; College of Science, Health, Engineering and Education, Murdoch University, Perth, WA 6150, Australia; Mayo Clinic Depression Center, Department of Psychiatry and Psychology, Mayo Clinic, Rochester, MN 55905, USA

**Author notes:** Corresponding Author Associate Professor Jennifer Rodger, M317, The University of Western Australia, 35 Stirling Highway, Crawley, Western Australia 6009.

## Abstract

Prior research suggests that the neurobiological underpinnings of depression include disruptions in functional connectivity, neurometabolite levels, and hippocampal volume. This study examined the validity of a chronic restraint stress (CRS) paradigm in male Sprague Dawley rats for the study of depression using longitudinal behavioural tests and multiple 9.4 T MRI modalities (resting-state functional MRI, proton magnetic resonance spectroscopy, and volumetric studies). In the CRS protocol, rats were placed in individual transparent tubes for 2.5 h daily over 13 days. Elevated plus-maze test (EPM) and forced swim test (FST) confirmed the presence of anxiety-like and depression-like behaviours respectively post-restraint. Brain changes were also detected by MR. The rs-fMRI data revealed hypoconnectivity within the salience and interoceptive networks and hyperconnectivity of several brain regions to the cingulate cortex. The ^1^H-MRS data revealed decreased sensorimotor cortical glutamate, glutamine and combined glutamate-glutamine levels. Volumetric analysis of T2-weighted images revealed decreased hippocampal volume, which was also correlated with salience network connectivity. Depression-like behaviours were correlated with salience and interoceptive network connectivity, glutamate and combined glutamate-glutamine levels and hippocampal volume. Anxiety-like behaviours were correlated with both hippocampus connectivity and interoceptive network connectivity. The present findings identify significant changes in brain connectivity, neurometabolites and structure that are correlated with abnormal behaviour in CRS rats. Importantly, these changes parallel those found in human depression, suggesting that the CRS rodent model has utility for translational studies and novel intervention development for depression.

## Introduction

Major depressive disorder is a debilitating neuropsychiatric disease with significant morbidity and mortality. The diagnosis of depression in humans is based on persistent negative mood, clinical symptoms, and behavioural changes. Diagnosing depression based solely on clinical features leads to suboptimal outcomes in research studies and clinical practice. Considerable effort has been focused on addressing the biological heterogeneity of depression with biomarkers, including magnetic resonance imaging (MRI) techniques. For example, resting-state functional MRI (rs-fMRI) studies of depression demonstrate changes in resting-state networks (RSNs) such as the salience network, interoceptive network and default mode network (DMN), involved in processing emotions and sensory stimuli and regulating the internal state (Paulus and Stein, 2010; Mulders *et al.*, 2015). These alterations in functional connectivity within RSNs are associated with neurometabolite (e.g., glutamine, Gln; glutamate, Glu; γ-aminobutyric acid, GABA) imbalances in depression, measured non-invasively using proton magnetic resonance spectroscopy (^1^H-MRS) (Lener *et al.*, 2017). Additionally, human MRI studies reproducibly detect reduced hippocampal volumes in patients with depression compared to age-matched healthy controls (Videbech and Ravnkilde, 2004).

Although significant progress has been made in understanding the mechanisms underpinning major depressive disorder, the causality of neuroimaging findings is difficult to infer. For example, the state vs trait nature of human imaging findings are often uncertain and difficult to study (e.g., Sheline, 2011; Brown *et al.*, 2014). In contrast, temporal relationships between biological findings and depression-like behaviours can be studied in animal models. Moreover, MRI-based techniques can be used to investigate the same biological changes in humans and animals, allowing direct comparison of validated outcome measures. Furthermore, combining repeated behavioural and MRI measures within the same animals allows the exploration of correlation between those measures. Therefore, animal models are an indispensable tool for studying aetiology, progress and treatment of depression.

Chronic restraint stress (CRS) in Sprague Dawley rats has been shown to elicit behavioural, genetic, protein, brain functional connectivity and hippocampal volume changes similar to those in patients with depression. However, no study to date has examined the association between multimodal MRI measures and behavioural changes within the same animals in the CRS model of depression (Lee *et al.*, 2009; Henckens *et al.*, 2015; Wang *et al.*, 2017; Alemu *et al.*, 2019). This model involves restraining animals’ movements for at least two hours a day for several days (Wang *et al.*, 2017); the continuous and predictable stress is designed to mimic every-day human stress, such as daily repetition of a stressful job and familial stresses. Our study aimed to investigate the relationship between neurobiological and behavioural changes in the CRS rat model by performing multimodal MRI (rs-fMRI, ^1^H-MRS, and structural MRI) and behavioural tests on the same animals before and after induction of the model.

## Materials and Methods

### Ethics statement

Experimental procedures were approved by the University of Western Australia (UWA) Animal Ethics Committee (RA/3/100/1640) and conducted in accordance with the *National Health and Medical Research Council Australian code* for the care and use of animals for scientific purposes.

### Animals

Adult male Sprague Dawley rats (n=30; 150-200 g; 6-7 weeks old) from the Animal Resources Centre (Canning Vale, WA, Australia) were housed under temperature-controlled conditions on a 12-hour light-dark cycle. Food and water were freely provided, except during the chronic restraint stress (CRS) procedure and fasting before the sucrose preference test. All rats were allowed to acclimatize to their new environment for one week following their arrival. Behavioural tests and MRI scans were carried out at baseline and after the last restraint procedure. A control group of (n=5) animals underwent all procedures except CRS.

### Chronic restraint stress procedure

The CRS procedure was carried out on a bench located on the opposite side of the animal holding room. Each session was carried out between 12:30 and 15:30 in effort to avoid effects associated with the circadian rhythm. In brief, rats were weighed and placed in a transparent tube (size of the tube depending on weight of animal, see Table 1) for 2.5 h a day for 13 consecutive days (Ulloa *et al.*, 2010). The length of each restraint was adjusted to limit limb movements using tail gates. Following CRS, rats were returned to their home cages. Healthy control animals were not restrained and remained in their home cages.

**Table 1.** Weight of animals and size of restraints.

| Body weight of animal (g) | Diameter of restraint (cm) | Maximum length of restraint (cm) |
| --- | --- | --- |
| < 255 | 5 | 19 |
| 255-300 | 5 | 23 |
| > 300 | 6 | 21 |

### Behavioural tests

Behavioural tests were carried out over a period of 3 days (Fig. 1). On the first day, the animals were subjected to elevated plus-maze (EPM) test (Walf and Frye, 2007). After EPM, the animals were habituated to single housing and 1% sucrose solution as described below and deprived of food and water overnight. The next day, the sucrose preference test (SPT) (Willner *et al.*, 1987) was carried out and on the third day, the animals underwent the forced swim test (FST) (Slattery and Cryan, 2012). All behavioural testing occurred between 08:30 and 11:00. The full behavioural dataset can be obtained from the corresponding author upon request.

**Figure 1.**
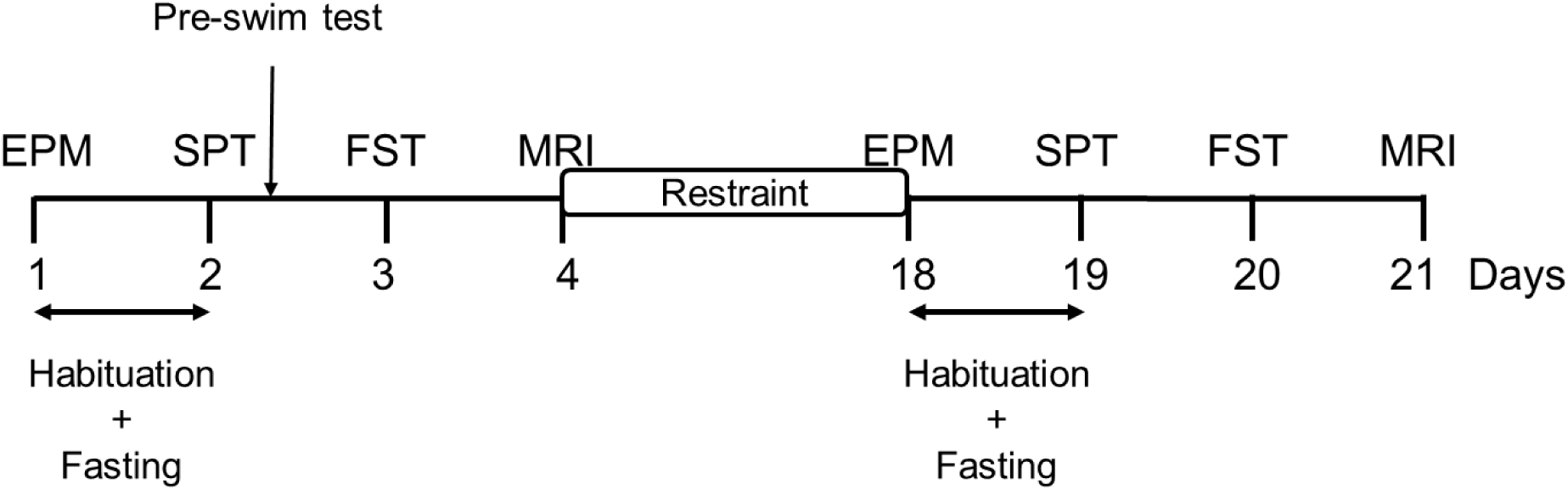
Experimental timeline. The experiment consisted of an initial one-week period of habituation upon arrival of the animal, after which the rats underwent elevated plus-maze (EPM) test (Day 1). Following EPM, animals were habituated to single housing and sucrose solution for 8 h and deprived of food and water overnight (Day 1-2). Sucrose preference test (SPT) was carried out the next day (Day 2), followed by a pre-swim test. Forced swim test (FST) was carried out on Day 3 and MRI on Day 4. The animals then underwent chronic restraint stress for 2.5 h daily for 13 consecutive days. The day after the end of CRS, animals underwent behavioural tests (Day 18-20) and MRI (Day 21) in the same order without a pre-swim test.

### Elevated plus maze

EPM was carried out as detailed in Walf and Frye (2007). The apparatus consisted of two open arms (without walls or railing) and two closed arms, crossed in the middle perpendicularly to each other, and a centre area (10 cm x 10 cm). Each arm was 50 cm long and 10 cm wide and the enclosed arms had 40 cm high walls. Each arm of the maze was attached to sturdy legs, such that it was elevated 60 cm off the floor. The maze was placed in a way to ensure similar levels of illumination on both open and closed arms. One animal was tested at a time and after each trial, all arms and the centre area were wiped with 70% ethanol to remove olfactory cues. The animal was placed in the centre of the maze facing the same open arm, away from the experimenter. The animal was allowed to move freely in the maze for 5 min and the whole procedure was video recorded from approximately 120 cm above the platform using a GoPro HERO7. The experimenters stayed in the room during the procedure, but unnecessary movements and noise were minimised.

EPM data was analysed manually by researchers blinded to experimental group and timepoint. The number of entries and time spent in closed and open arms were measured. Additionally, the number and duration of rearing and grooming were also measured to investigate anxiety-related behaviours (Walf and Frye, 2007). The number of entries and time spent in the centre of the maze and behaviours such as head shaking, head dips and stretching were not considered. One animal fell off the open arm during baseline testing and replaced onto the maze to continue the whole 5 min testing but the data was excluded from the analyses (Walf and Frye, 2007).

### Sucrose preference test

Immediately following EPM, animals were habituated to single housing and to sucrose solution (Fig. 1). Animals were placed in individual cages with *ad libitum* access to food pellets and two 600 ml bottles, one bottle containing fresh 1% sucrose solution and the other containing tap water. The animals were trained to this condition for 8 h. Rats were given a free choice between the two bottles and the position of the bottles was switched 4 h after the start of single housing to prevent side preference in drinking behaviour. Overnight food and water deprivation was applied at the end of the 8 h habituation for up to 16 h.

The next day, SPT was performed according to a previous study with some modifications (Willner *et al.*, 1987). Water and sucrose solution bottles were weighed, labelled and placed in corresponding cages. The position of the bottles was switched 30 min after the start of the SPT. 30 min later, the bottles were removed and re-weighed, and the animals were re-housed in their original cages. Sucrose preference was calculated as a percentage of the total amount of liquid ingested (sucrose preference = sucrose consumption (g) / [sucrose consumption (g) + water consumption (g)]).

### Forced swim test

FST was carried out as detailed in Slattery and Cryan (2012). Briefly, 20 L white opaque plastic buckets (41 cm high, 28 cm wide) were filled up to a depth of 30 cm with water at 23-25 °C. At this depth, the rats could not touch the bottom of the bucket with their tails or hind limbs. Up to four buckets were used at a time and the buckets were emptied, cleaned and refilled between animals. At baseline, 24 h before the FST session (on SPT day), rats were exposed to a pre-swim test for 10 min by placing them in the water-filled buckets (Slattery and Cryan, 2012). The next day, and at the end of the restraint period rats underwent 6 min of FST (Fig. 1) and the procedure was video recorded from approximately 50 cm above the buckets using a GoPro HERO7.

FST data was analysed manually by researchers blinded to experimental group and timepoint using a time-sampling technique (Slattery and Cryan, 2012). The first 5 min of the video recording was split into 5-s intervals and the predominant behaviour in each 5-s period was rated. The following escape behaviours were scored: (i) swimming, with horizontal movements throughout the bucket including crossing into another quadrant and diving; (ii) climbing, with upward movements of the forepaws along the side of bucket; (iii) immobility, with minimal movements necessary to keep their head above water; and (iv) latency, defined as the time taken to exhibit the first immobility behaviour. Grooming, head shaking and number of fecal boli were not considered. Trials during which the animals managed to escape more than once or were floating horizontally for the duration of the test (with most of their body being completely dry at the end) were excluded from the analyses.

### MRI data acquisition

#### Animal preparation

MRI data was acquired the day after the FST. The rat was pre-anaesthetised in an induction chamber (4% isoflurane in medical air, 2 L/min). Once fully anaesthetised, the animal was transferred to a heated imaging cradle and anaesthesia was maintained with a nose cone (2% isoflurane in medical air, 1 L/min). Body temperature and respiratory rate were monitored using a PC-SAM Small Animal Monitor (SA Instruments Inc., 1030 System). An MR-compatible computer feedback heating blanket was used for maintaining animal body temperature at 37°C (± 0.5°C). A 25G butterfly catheter was implanted subcutaneously in the left flank of the animal to deliver a 0.05-0.1 mg/kg bolus injection and continuous infusion of medetomidine at 0.15 mg/kg/h using an infusion pump. Once the animal’s breathing rate dropped to 50 breaths/min, isoflurane was gradually reduced to 0.5-0.75%. These anaesthetic doses were empirically determined to ensure the respiratory rate of the animals was between 50-80 breaths/min. rs-fMRI scans were started only after the isoflurane concentration had been reduced for at least 15 min and the physiology of the animal was stable during that time. After the MRI procedure, medetomidine was antagonized by an injection of atipamezole (0.1 mg/kg).

#### MRI acquisition parameters

All MR images were acquired with a Bruker Biospec 94/30 small animal MRI system operating at 9.4 T (400 MHz, H-1), with an Avance III HD console, BGA-12SHP imaging gradients, a 72 mm (inner diameter) volume transmit coil and a rat brain surface quadrature receive coil using the imaging protocol as described in detail in Seewoo *et al.* (2018; 2019). Briefly, the acquisition protocol included the following sequences: 1) multi-slice 2D RARE (rapid acquisition with relaxation enhancement) sequence for three T2-weighted anatomical scans (TR=2500 ms, TE=33 ms, matrix=280×280, pixel size=0.1×0.1 mm^2^, 21 coronal and axial slices, 20 sagittal slices, thickness=1 mm); 2) single-shot gradient echo EPI (TR=1500 ms, TE=11 ms, matrix=94×70, pixel size=0.3×0.3 mm^2^, 21 coronal slices, thickness=1 mm, flip angle=90°, 300 volumes, automatic ghost correction order = 1, receiver bandwidth = 300 kHz) for resting-state; and 3) point-resolved spectroscopy (PRESS) sequence with one 90° and two 180° pulses and water suppression for ^1^H-MRS (TE=16 ms, TR=2500 ms) with 64 averages with a 3.5×2×6 mm^3^ voxel placed over the left sensorimotor cortex (Fig. 2). ^1^H-MRS data was acquired from the left sensorimotor cortex only in order to facilitate comparisons with human depression studies, which mostly examine neurometabolite changes in the left hemisphere (Moriguchi *et al.*, 2019).

**Figure 2.**
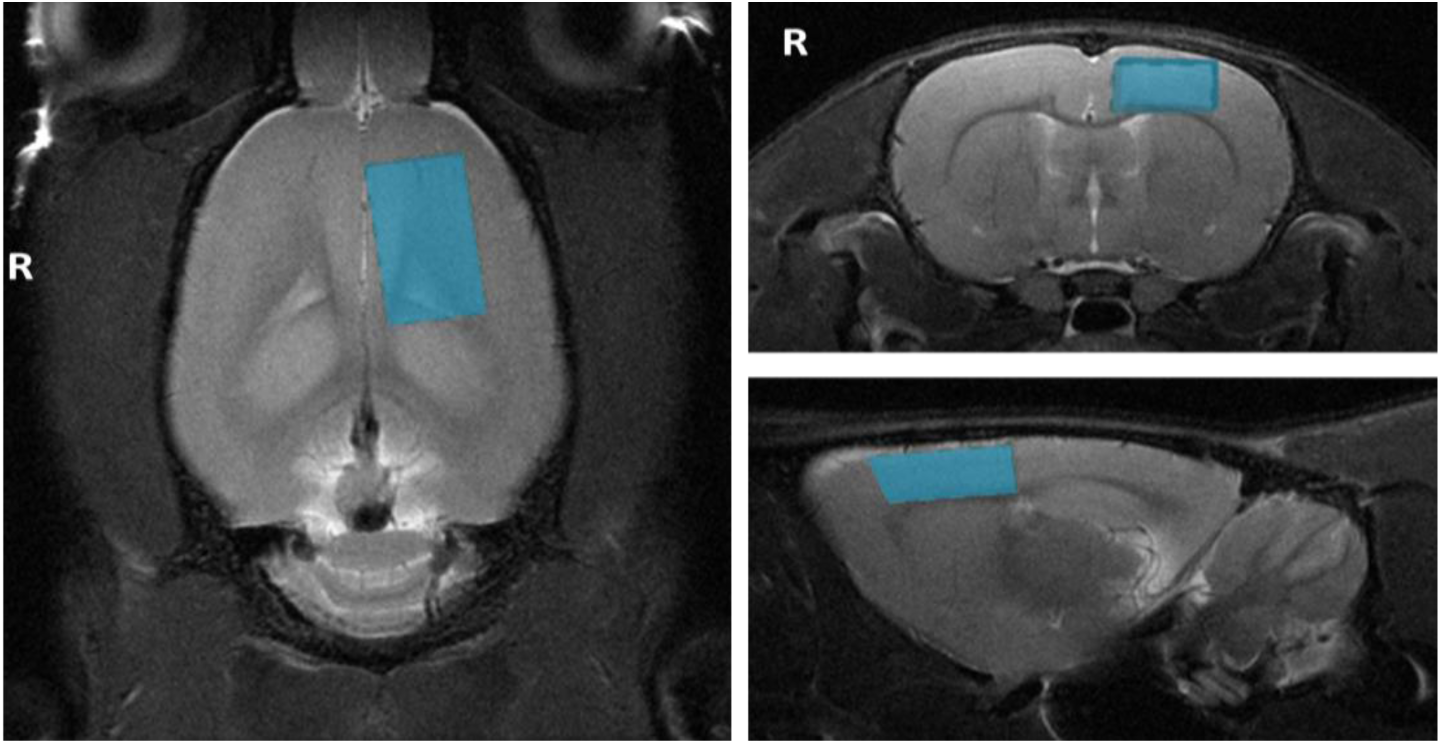
Voxel position for proton magnetic resonance spectroscopy. The figure shows the position of the voxel of interest (size of 3.5 mm x 2 mm x 6 mm) on the left sensorimotor cortex on T2-weighted images for proton magnetic resonance spectroscopy.

### Statistical analyses

Shapiro-Wilk normality tests were conducted before each analysis (except rs-fMRI), followed by a t-test or Wilcoxon test. Unless otherwise stated, all results are expressed as mean ± SEM and *p* < 0.05 was considered significant. Table 2 lists the specifications of each test.

**Table 2.**
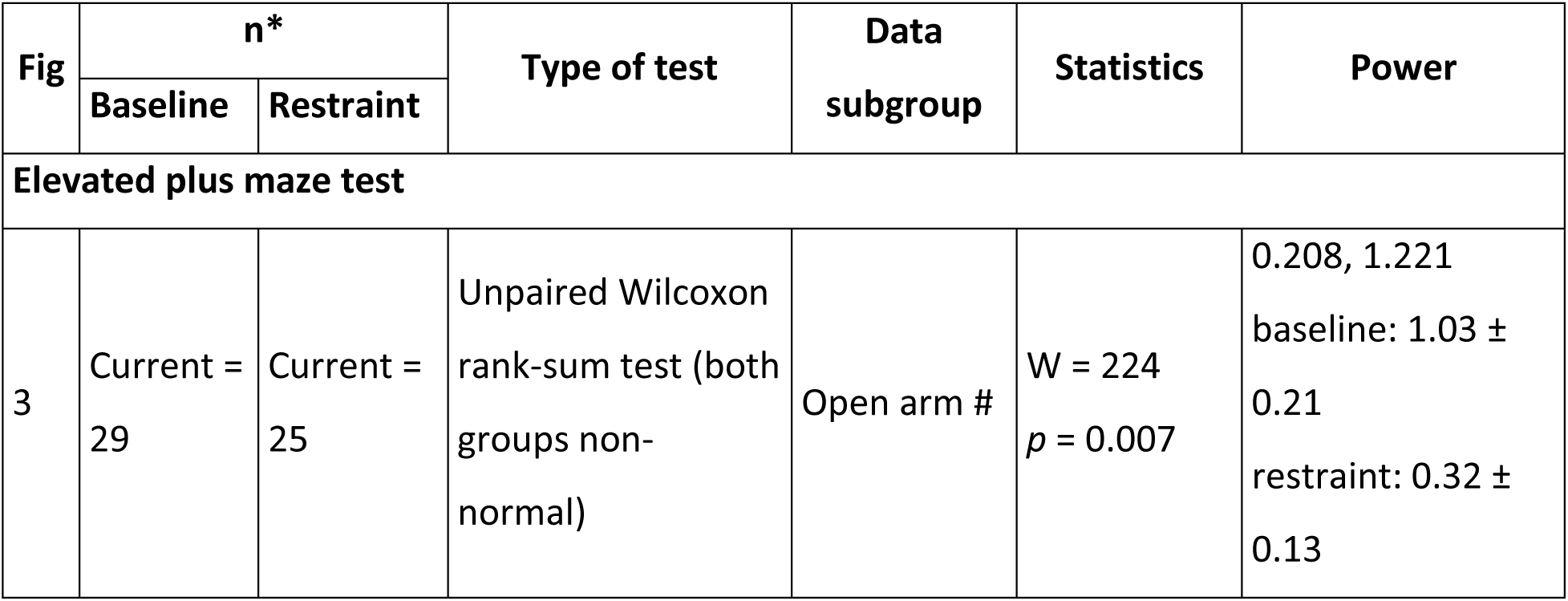
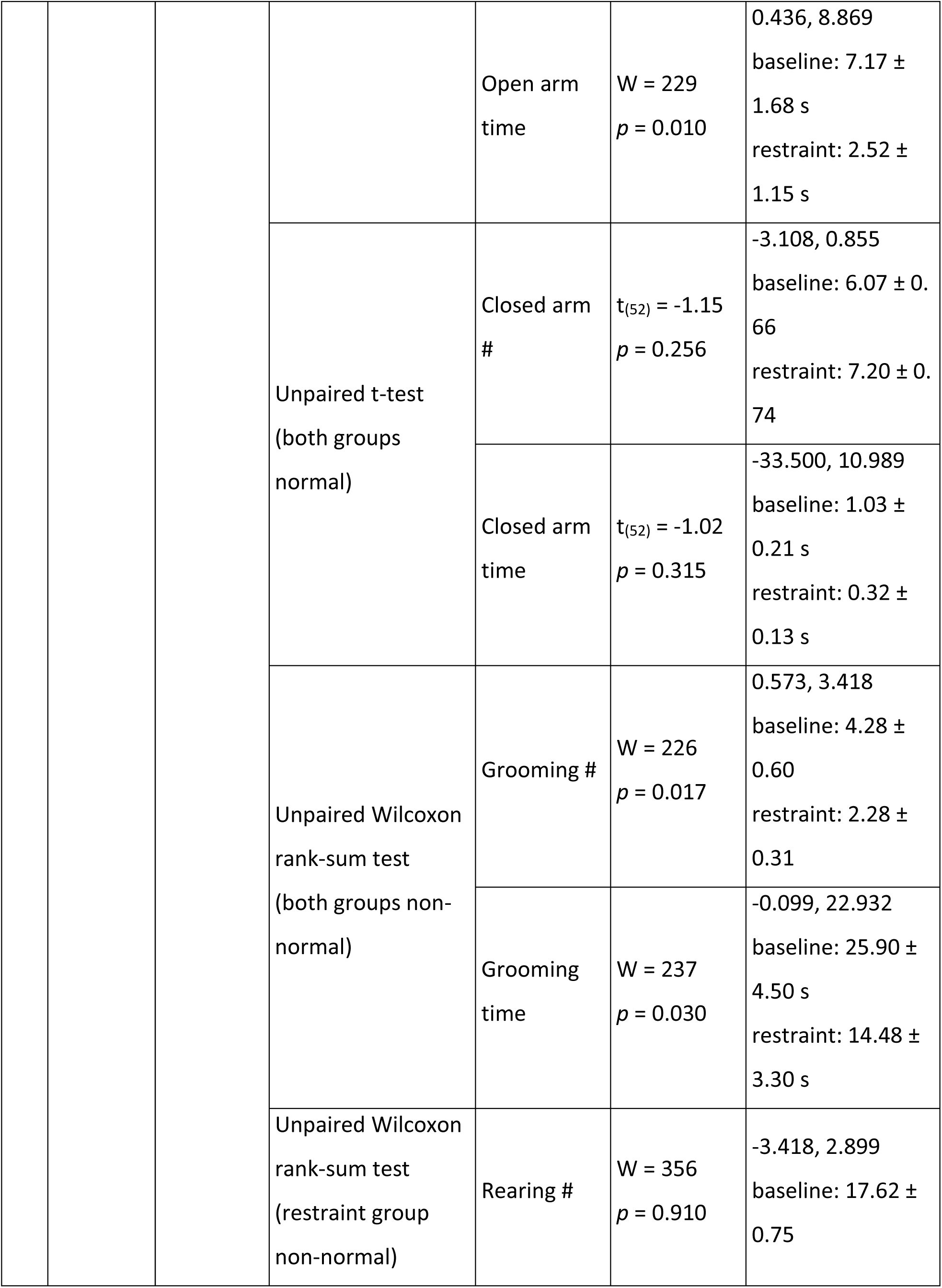
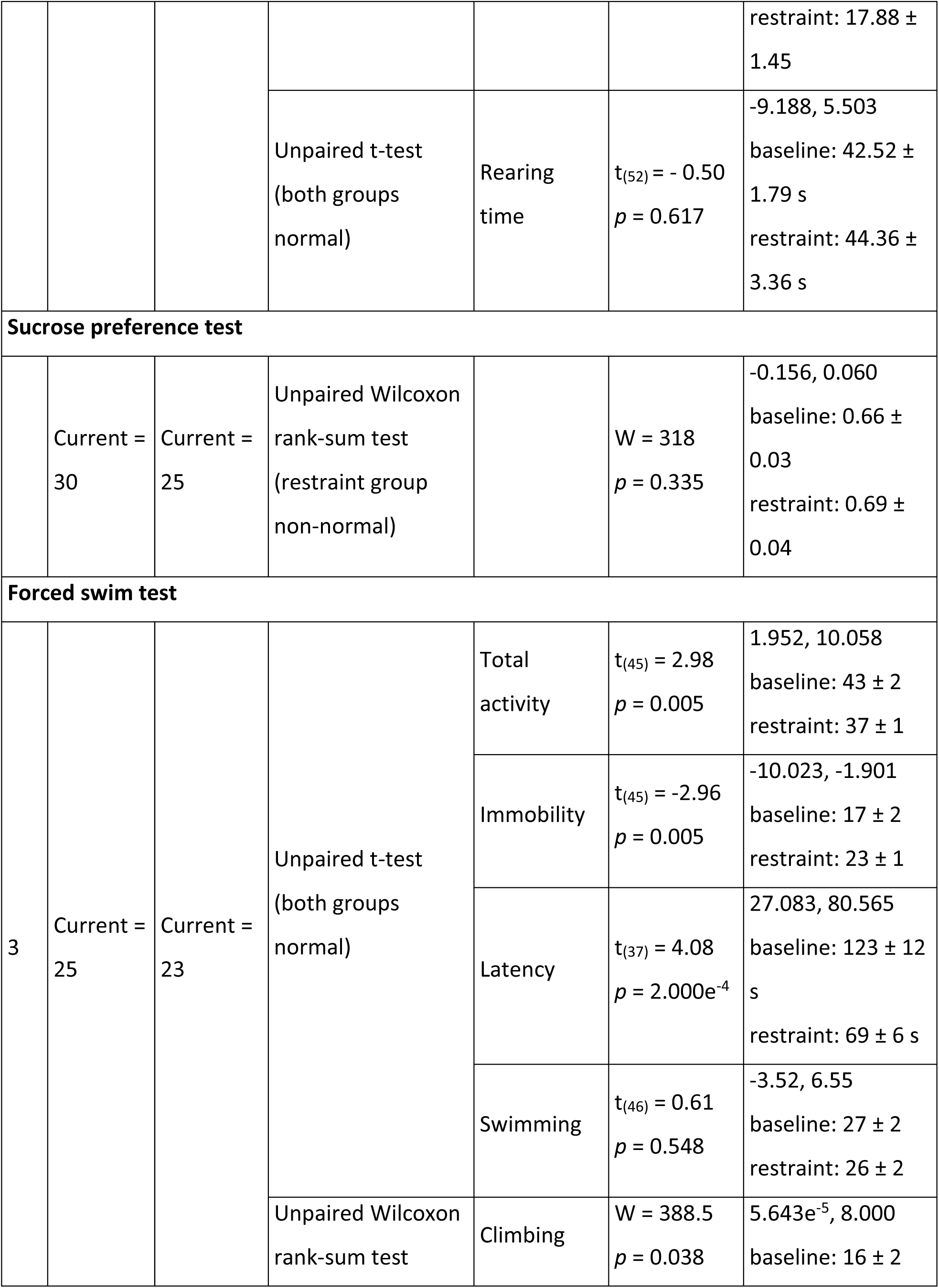
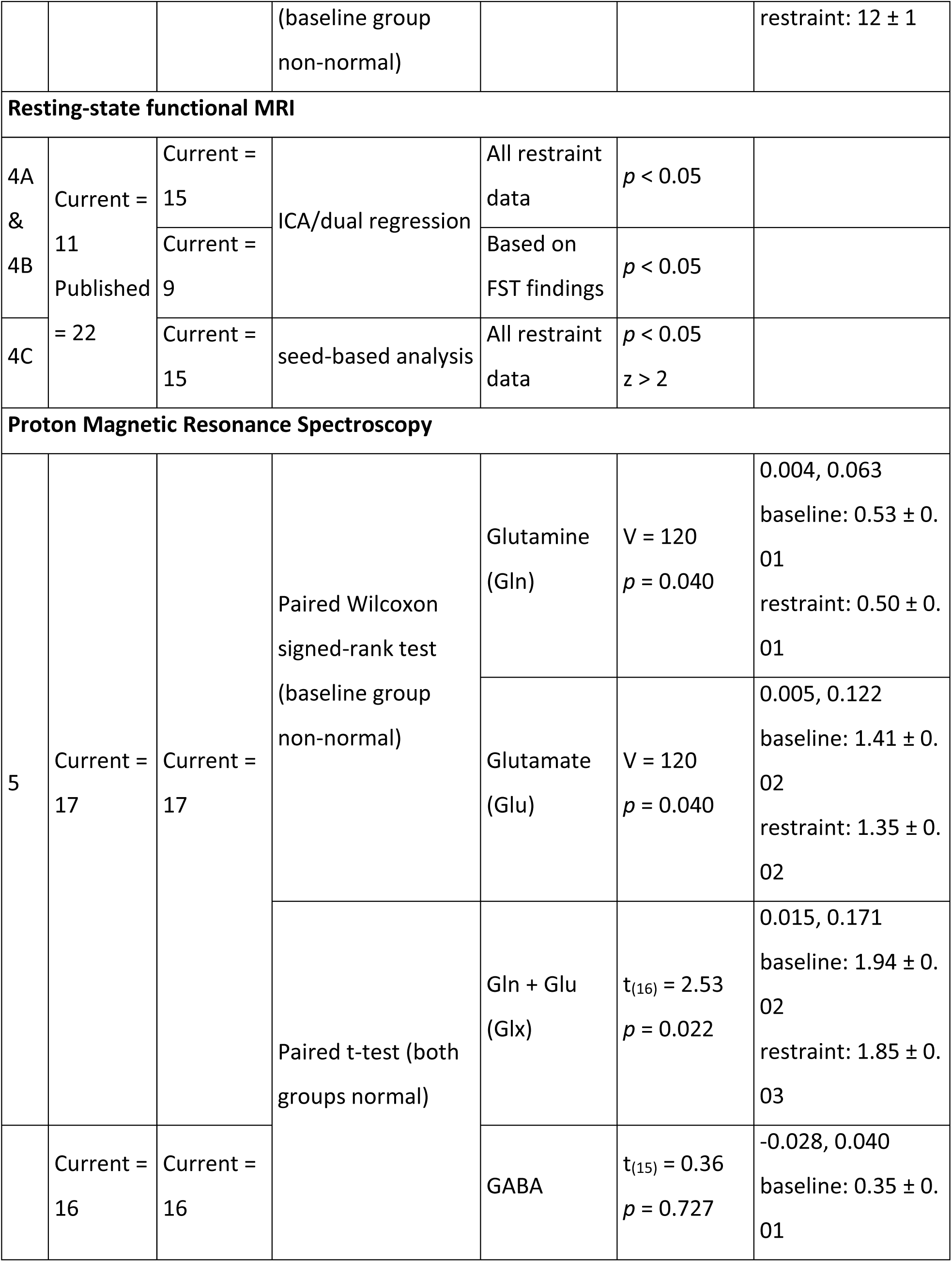
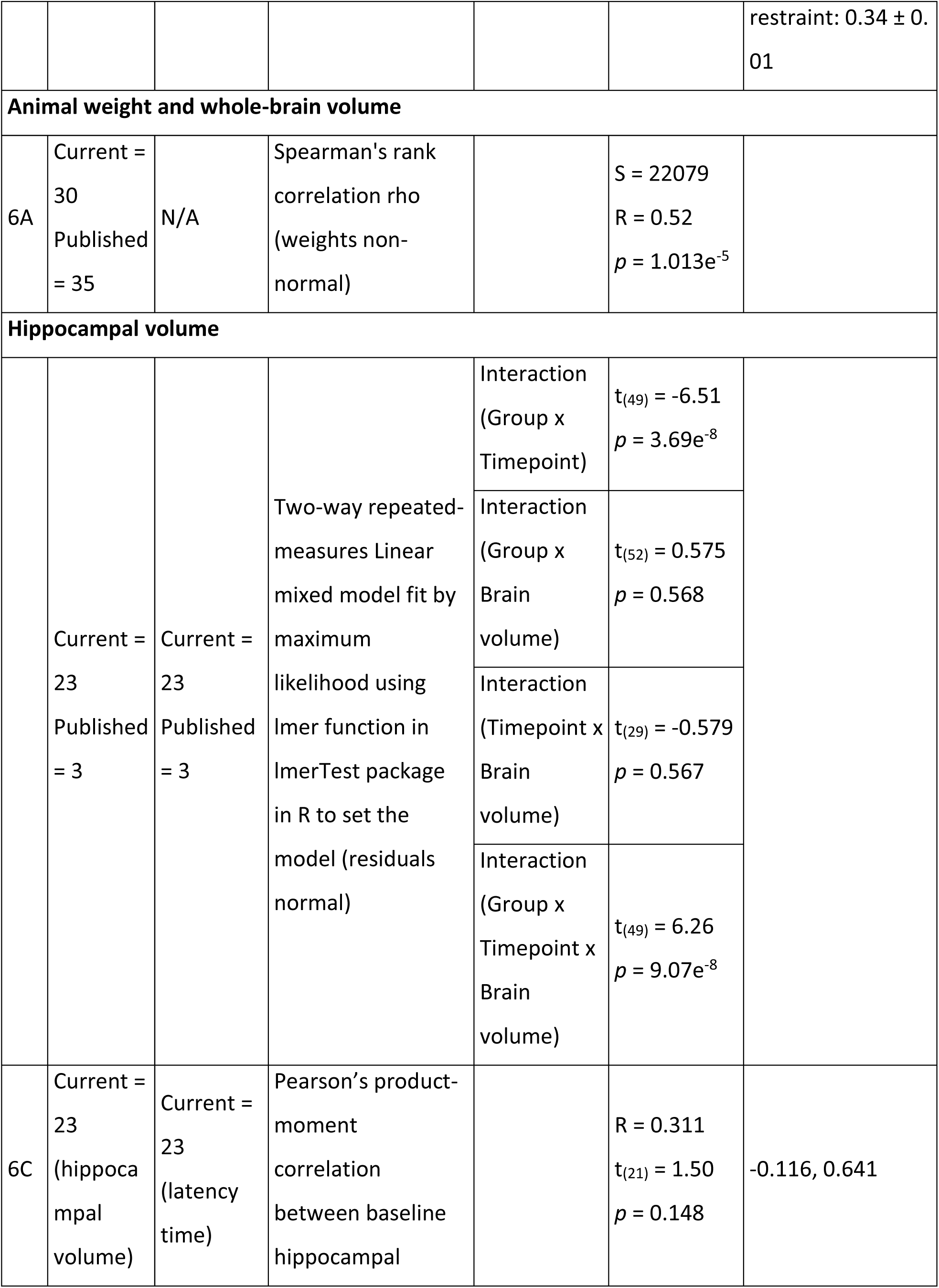
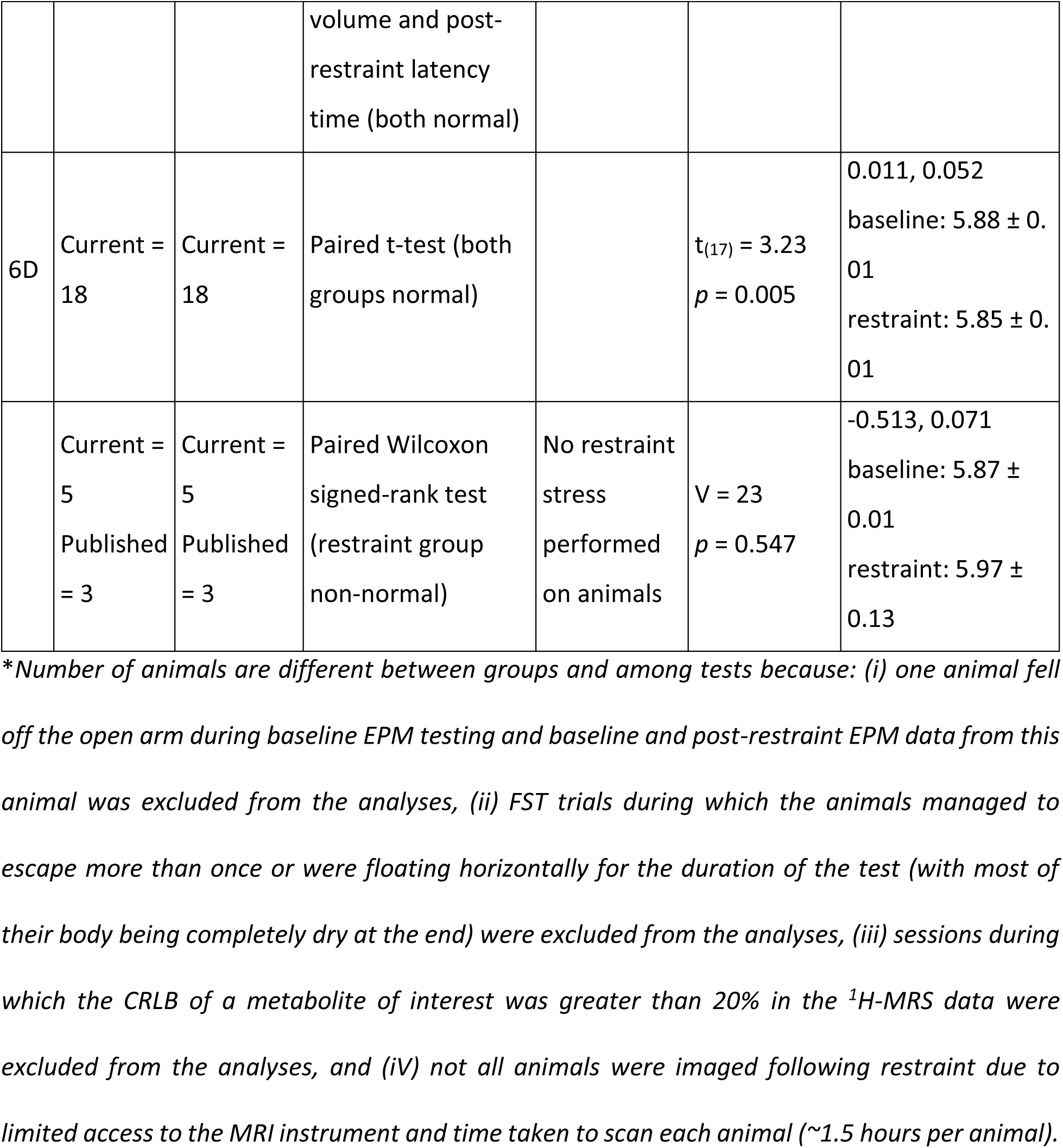
Statistical table indicating the results of all analyses. Each analysis includes a letter indicator linking the test in the table to the analysis in the text. The link to the corresponding figure, if any, is indicated under “Fig.”. Under “n”, the number of animals used from “current” cohort and from the “published” data from Seewoo et al. (2019) in each group (baseline and restraint) is indicated. Shapiro-Wilk normality test was used to determine the distribution of each group, with p > 0.1 indicating normal distribution and the results are mentioned under “Type of test” (normal or non-normal distribution). The critical value, degrees of freedom and p values are listed for each test under “Statistics” and the confidence intervals of the tests are under “Power”.

#### Behavioural data

EPM and FST videos were scored blind by two trained observers (BJS & LAH) to establish the most reliable measures. Based on their low inter-individual scorer variability (<12%), exploration (into and within closed and open arms separately), grooming and rearing for EPM, and swimming and climbing (separately and combined as “total activity”), immobility, and latency to first immobility behaviour for FST were selected for statistical analysis. Sucrose preference (%) was calculated as sucrose consumption (g)/[sucrose consumption (g) + water consumption (g)]. Behavioural data were analysed with unpaired Wilcoxon signed-rank tests or unpaired t-tests (Table 2).

#### Rs-fMRI data

To maximise the use of collected data, rs-fMRI data and T2-weighted images collected using the same acquisition and anaesthesia protocols from a previous study (Seewoo *et al.*, 2019) in adult (6–8 weeks old, 150–250 g) male Sprague Dawley rats were also included in the analyses for baseline and control groups (Table 2). These animals did not undergo any behavioural testing or intervention before the acquisition of MRI data. All rs-fMRI data was pre-processed and analysed in the same way. Pre-processing of data included: (i) export into DICOM format from ParaVision 6.0.1 (Bidgood Jr *et al.*, 1997); (ii) conversion into NifTI using the dcm2niix converter (64-bit Linux version 5 May 2016) (Rorden *et al.*, 2007); (iii) reorienting the brain into left-anterior-superior (LAS) axes (radiological view); and (iv) skull-stripping using the qimask utility from QUIT (QUantitative Imaging Tools) (Wood, 2018). The voxel sizes were then upscaled by a factor of 10 (Tambalo *et al.*, 2015).

All further pre-processing and analyses were performed using FSL v5.0.10 (Functional MRI of the Brain (FMRIB) Software Library) (Jenkinson *et al.*, 2012). Single-session independent component analysis (ICA) as implemented in FSL/MELODIC (Multivariate Exploratory Linear Decomposition into Independent Components) (Beckmann *et al.*, 2005) was used to de-noise the data (detailed in Seewoo *et al.*, 2018). The de-noised rs-fMRI images were then co-registered to their respective T2-weighted coronal images using six-parameter rigid body registration using FSL/FLIRT (Linear Image Registration Tool) (Jenkinson and Smith, 2001; Jenkinson *et al.*, 2002) and normalised to a Sprague-Dawley brain atlas (Papp *et al.*, 2014; Kjonigsen *et al.*, 2015; Sergejeva *et al.*, 2015) with nine degrees of freedom ‘traditional’ registration. The atlas was first down-sampled by a factor of eight to better match the voxel size of the 4D functional data. All subsequent analyses were conducted in the atlas standard space.

Multi-subject temporal concatenation group-ICA and FSL dual regression analysis were used to determine group differences (baseline, n=33; restraint, n=15), controlling for family-wise error (FWE) and using a threshold-free cluster enhanced (TFCE) technique to control for multiple comparisons. The resulting statistical maps were thresholded to p < 0.05.

To investigate the correlation between strong depression-like behaviours on functional connectivity, dual regression was also carried out using a subset of animals in the restraint group. FST measures (immobility, swimming and climbing scores and latency time) were extracted for the 15 animals scanned following restraint and sorted in order of greatest change in each FST measure. Animals were scored according to their position on the list (1 to 15). Nine of 15 animals had consistently high scores and were used in the analysis as they exhibited the greatest change in overall behavioural outcomes in FST.

For seed-based analysis, the atlas mask for cingulate cortex was transformed to each individual animal’s functional space. The region of interest (ROI) masks (within the individual functional space) were used to extract the timecourse from the ICA de-noised data. The timecourses were used in a first-level FSL/FEAT (FMRI Expert Analysis Tool Version 6.00) analysis to generate a whole-brain correlation map. Higher-level analysis was carried out using OLS (ordinary least squares) simple mixed-effects (Beckmann *et al.*, 2003; Woolrich *et al.*, 2004; Woolrich, 2008) in atlas space (baseline, n=33; restraint, n=15). Z (Gaussianised T/F) statistic images were thresholded non-parametrically using clusters determined by Z > 2 and a (corrected) cluster significance threshold of *p* = 0.05 (Worsley, 2001).

#### ^1^H-MRS

^1^H-MRS data were analysed in LCModel (“Linear Combination of Model spectra” version 6.3-1L) (Provencher, 2001) using a set of simulated basis set provided by the software vendor. For data quality control, the linewidth (full width at half-maximum, FWHM) for each scan was calculated for the N-acetylaspartate+N-acetyl-aspartyl-glutamate (NAA + NAAG) resonance at 2.01 ppm and the intensity of this resonance relative to the residual intensity was obtained (signal-to-noise ratio, SNR). Mean (± SE) FWHM and SNR were 10.6 (± 0.4) Hz and 12.1 (± 0.4) respectively for the baseline group (n=17), and 14.6 (± 0.8) Hz and 9.8 (± 0.3) respectively for the restraint group (n=17). Individual metabolite concentrations were computed using the unsuppressed reference water signal for each individual scan. Cramér-Rao lower bound (CRLB) values were calculated by LCModel and reported as percent standard deviation of each metabolite, as a measure of the reliability of the metabolite estimates.

The metabolites of interest were GABA (baseline CRLB: 12.7% ± 0.4, restraint CRLB: 14.5% ± 0.6) and Glu (baseline CRLB: 3.4% ± 0.1, restraint CRLB: 3.9% ± 0.1), the major neurotransmitters in the brain, as well as Gln (baseline CRLB: 9.7% ± 0.5, restraint CRLB: 11.4% ± 0.3), a neurotransmitter precursor, and combined glutamate-glutamine (Glx; baseline CRLB: 3.6% ± 0.1, restraint CRLB: 4.2% ± 0.1). To accurately extract the dominating metabolic changes observed before and after CRS, and to reduce systemic variations among studied animals, a relative quantification method, using an internal spectral reference was used. All ^1^H-MRS results presented here are expressed as a ratio to tCr (total creatine = Cr + PCr; baseline CRLB: 2.94% ± 0.06, restraint CRLB: 3.12% ± 0.08) spectral intensity, the simultaneously acquired internal reference peak (Block *et al.*, 2009; Walter *et al.*, 2009; Xu *et al.*, 2013). Paired Wilcoxon signed-rank test or paired t-test was used to test for changes in the metabolite ratios between baseline and restraint groups (Table 2).

#### Hippocampal volume

The three T2-weighted anatomical (coronal, sagittal and axial) data were pre-processed as above and then registered to the high-resolution atlas (no downsampling). Atlas masks for bilateral hippocampus and whole-brain were used to automatically extract their respective volumes from the three T2-weighted anatomical images (coronal, sagittal and axial). Hippocampal and whole-brain volumes from the three planes were averaged for each animal scan session. Spearman’s rank correlation method (n=65) was used to determine the correlation of whole-brain volumes with the weight of animals at baseline because the volumes of the whole brain and several brain regions are known to increase with age in rats until they are two months old (Mengler *et al.*, 2014). Using ‘lmer’ function from ‘lmerTest’ package in RStudio 3.5.2, the interaction effects of group (with CRS vs healthy), timepoint (baseline vs restraint) and whole-brain volume on raw hippocampal volumes were determined. Residuals from the model were normally distributed. Hippocampal volume was normalised to the whole-brain volume (% whole brain volume) to adjust for differences in head size (Welniak–Kaminska *et al.*, 2019). Pearson’s correlation method was used to determine correlation between baseline percentage hippocampal volume and post-restraint latency to first immobility behaviour during FST. A paired t-test was used to test for differences in percentage hippocampal volumes between baseline and restraint groups (n=18/group; Table 2) and a paired Wilcoxon signed-rank test was used to test for changes in percentage hippocampal volumes in healthy animals over two weeks without CRS (n=8/group; Table 2).

#### Correlations

Data from animals which underwent imaging at both timepoints were used to correlate MRI measures to the behavioural measures. Using the ‘rcorr’ function from the ‘Hmisc’ package in RStudio 3.5.2, Pearson or Spearman correlations (depending on normality of data) between the following parameters were computed: open arm entries and the number of grooming behaviours from EPM data; climbing score, immobility score and latency time from FST data; connectivity (average z-scores) of the hippocampus, cingulate cortex, primary somatosensory cortex and salience network from the rs-fMRI data; Glu and Glx ratios from ^1^H-MRS data; and hippocampal volume.

## Results

### Increase in anxiety and depression-like behaviours

In the elevated plus-maze (EPM) test, there was a significant decrease in the number of entries and exploration time in the open arms, but not in the closed arms of the maze (Fig. 3A, Table 2) following chronic restraint stress (CRS). The number of times rats demonstrated grooming behaviours and time spent grooming also significantly decreased following CRS, while the number of rearing behaviours and time spent rearing remained similar (Fig. 3B, Table 2).

**Figure 3.**
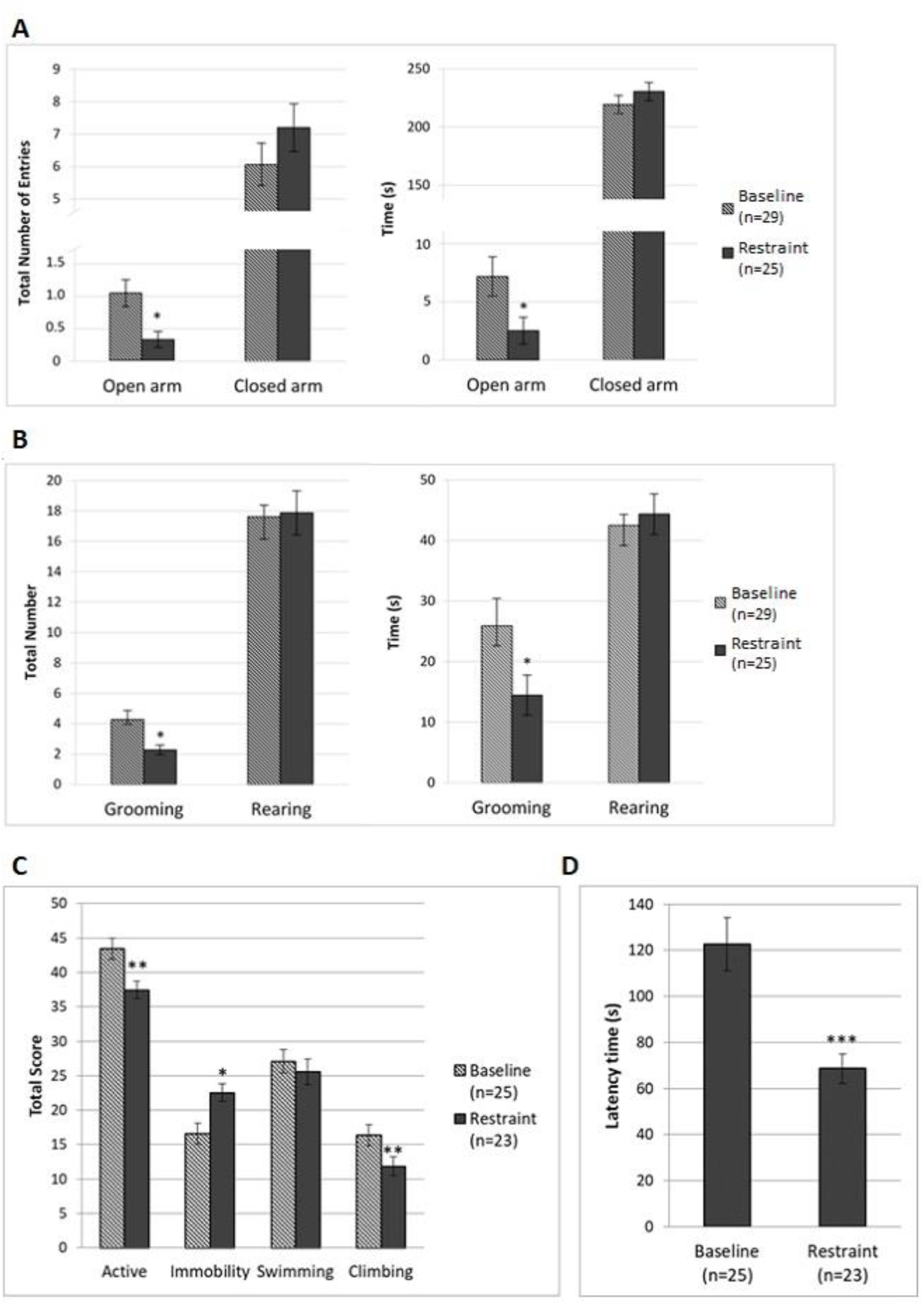
Effect of chronic restraint stress on behaviours displayed during elevated plus maze (A-B) and during forced swim test (C-D). For elevated plus-maze test, exploration (open and closed arm entries and time spent) and stress-related behaviours (grooming and rearing) were measured for 5 min (A-B; baseline, n=29; restraint, n=25) and for forced swim test, active behaviours (climbing and swimming), immobility and latency to first immobility behaviour were measured for 5 min (C-D; baseline, n=25; restraint, n=23). Data represent mean ± SEM. (A) Total number of entries and time spent in open arms decreased after 13 days of chronic restraint stress. (B) Number of grooming behaviours and time spent grooming decreased following restraint. (C) Scores for active behaviours decreased and immobility behaviour increased after 13 days of chronic restraint stress. When active behaviour scores were split into swimming and climbing, there was no change in swimming scores but a decrease in climbing scores. (D) Time to first immobility behaviour (known as latency time) decreased after restraint. Comparisons were made by an unpaired t-test. *p < 0.05, **p < 0.01, ***p < 0.001

An unpaired Wilcoxon signed-rank test on the sucrose preference test (SPT) data showed no significant changes in sucrose preference following CRS (Table 2).

For the forced swim test (FST), animals had similar scores for immobility and climbing behaviours at baseline, displaying total active behaviours (climbing plus swimming) for approximately 72% of the time (Fig. 3C, Table 2). After restraint, total activity decreased significantly compared to baseline while immobility increased and latency to first immobility behaviour decreased (Fig. 3D, Table 2). Additionally, when scores for both active behaviours (climbing and swimming) were split, the decrease in climbing behaviours following CRS was statistically significant, but not for swimming (Fig. 3C, Table 2).

### Changes in resting-state functional connectivity

The interoceptive (Becerra *et al.*, 2011; Seewoo *et al.*, 2019) and salience (Bajic *et al.*, 2016; Seewoo *et al.*, 2019) networks were identified from baseline rs-fMRI data and used in dual regression analysis for detecting functional connectivity differences induced by CRS (Fig. 4). Dual regression analysis revealed a significant decrease in connectivity of the bilateral somatosensory cortex to the salience network (Fig. 4A) and of the right somatosensory cortex to the interoceptive network (Fig. 4B). As a supplementary analysis, dual regression was carried out using a subset of the restraint group, which consisted of the nine animals exhibiting the greatest change in FST behavioural outcomes. A greater number of significant voxels with *p* < 0.05 (both networks) and lower p-values for changes in the salience network were obtained in the same brain regions (Fig. 4A-B; Table 3). Additionally, dual regression detected a significant decrease in connectivity of the right motor cortex and bilateral insular cortex to the salience network (Fig. 4A). Therefore, despite being a smaller group, the use of a subset of animals selected based on their FST performance resulted in increased sensitivity of the dual regression tool in detecting between-group differences. When rs-fMRI data of all animals were analysed using a seed-based analysis, a significantly greater functional connectivity of several brain regions to the cingulate cortex was detected in the restraint group (Fig. 4C). Specifically, hyperconnectivity was detected in the right retrosplenial cortex, visual cortex and inferior colliculus and in the bilateral thalamus, superior colliculus, dentate gyrus and Cornu Ammonis 3 (CA3).

**Table 3.** Summary of changes in functional connectivity within the interoceptive and salience networks when using all rs-fMRI data from post-restraint timepoint vs using a subset of animals showing greatest behavioural changes in forced swim test (FST).

| RSN | Contrast | Minimum p-value | Total number of significant voxels |
| --- | --- | --- | --- |
| Interoceptive network | Baseline > Restraint | 0.005 | 32 |
|  | Baseline > Restraint<sub>FST</sub> | 0.012 | 60 |
| Salience network | Baseline > Restraint | 0.029 | 11 |
|  | Baseline > Restraint<sub>FST</sub> | 0.002 | 127 |

**Figure 4.**
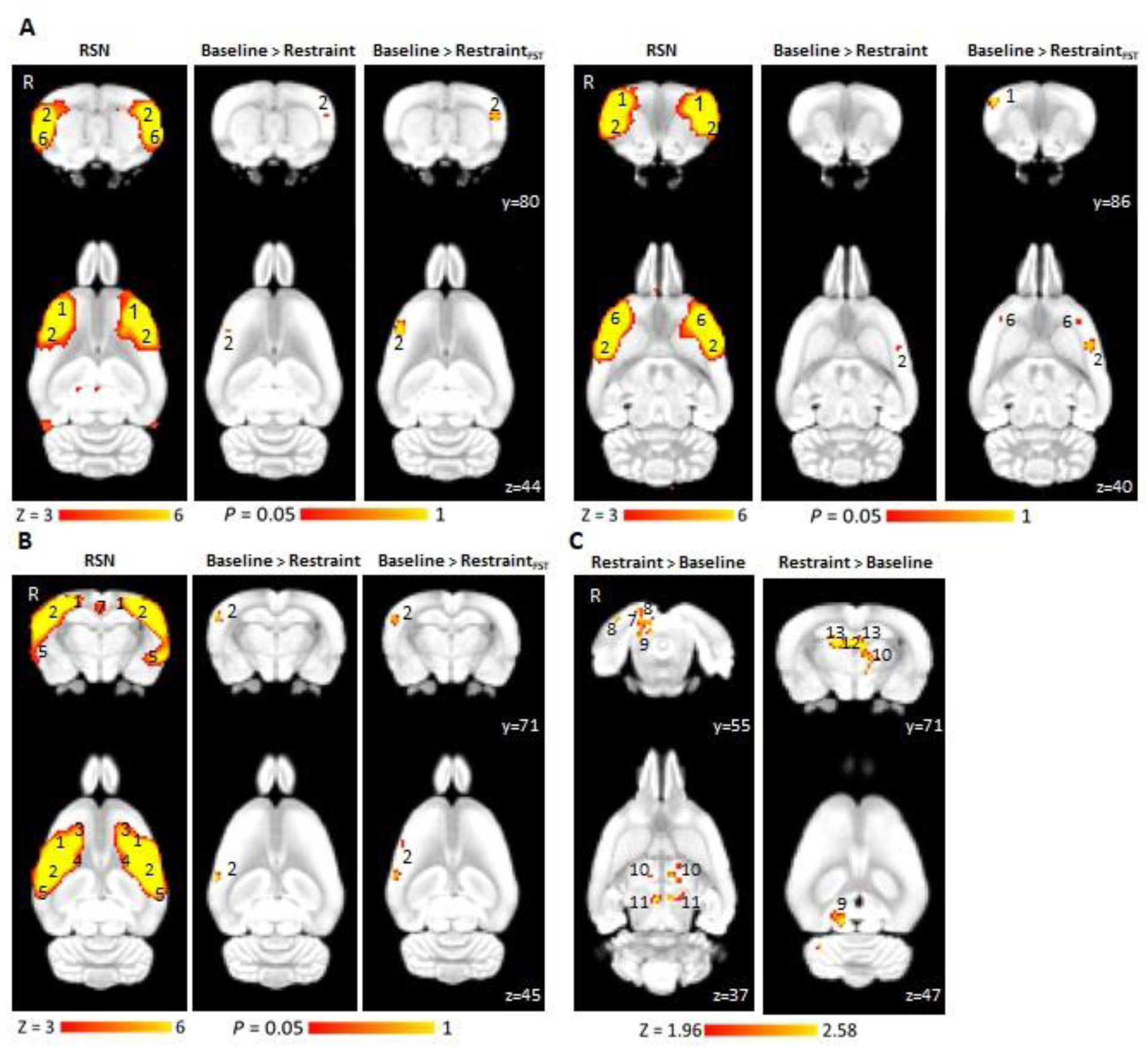
Decreased functional connectivity within the salience (A) and interoceptive networks (B) following chronic restraint stress as detected by dual regression and increased connectivity to the cingulate cortex as detected by seed-based analysis (C). The figure illustrates coronal and corresponding axial slices of spatial statistical colour-coded maps are overlaid on the rat brain atlas (down-sampled by a factor of eight). A and B show two RSNs (A, salience network and B, interoceptive network) identified in the baseline rs-fMRI scans of 6-7-week-old male Sprague Dawley rats under isoflurane-medetomidine anaesthesia. The RSN maps are represented as z-scores (n=33, thresholded at z > 3), with a higher z-score (yellow) representing a greater correlation between the time course of that voxel and the mean time course of the component. The changes in the functional connectivity within the two RSNs following 13 days of chronic restraint stress are represented as p-values (thresholded at *p* < 0.05; baseline, n=33; restraint, n=15; restraint based on FST result, n=9). C shows changes in the functional connectivity of the cingulate cortex between baseline and following 13 days of chronic restraint stress as spatial colour-coded Z (Gaussianised T/F) statistic images corrected for multiple comparisons at cluster level (thresholded at *p* < 0.05; baseline, n=33; restraint, n=15). R denotes right hemisphere. Significant clusters include various brain regions: 1, motor cortex; 2, somatosensory cortex; 3, frontal association cortex; 4, striatum/caudate putamen; 5, auditory cortex; 6, insular cortex; 7, retrosplenial cortex; 8, visual cortex; 9, inferior colliculus; 10, thalamus; 11, superior colliculus; 12, dentate gyrus; and 13, Cornu Ammonis 3 (CA3).

### Changes in neurometabolite levels as detected by ^1^H-MRS

The concentrations of the neurotransmitters GABA and Glu, the neurotransmitter precursor Gln, and combined glutamate-glutamine (Glx) were measured before and after CRS and were computed relative to tCr. Following restraint, rats had lower levels of Gln, Glu and Glx in the sensorimotor cortex (Fig. 5), while GABA levels were not significantly different from baseline (Table 2).

**Figure 5.**
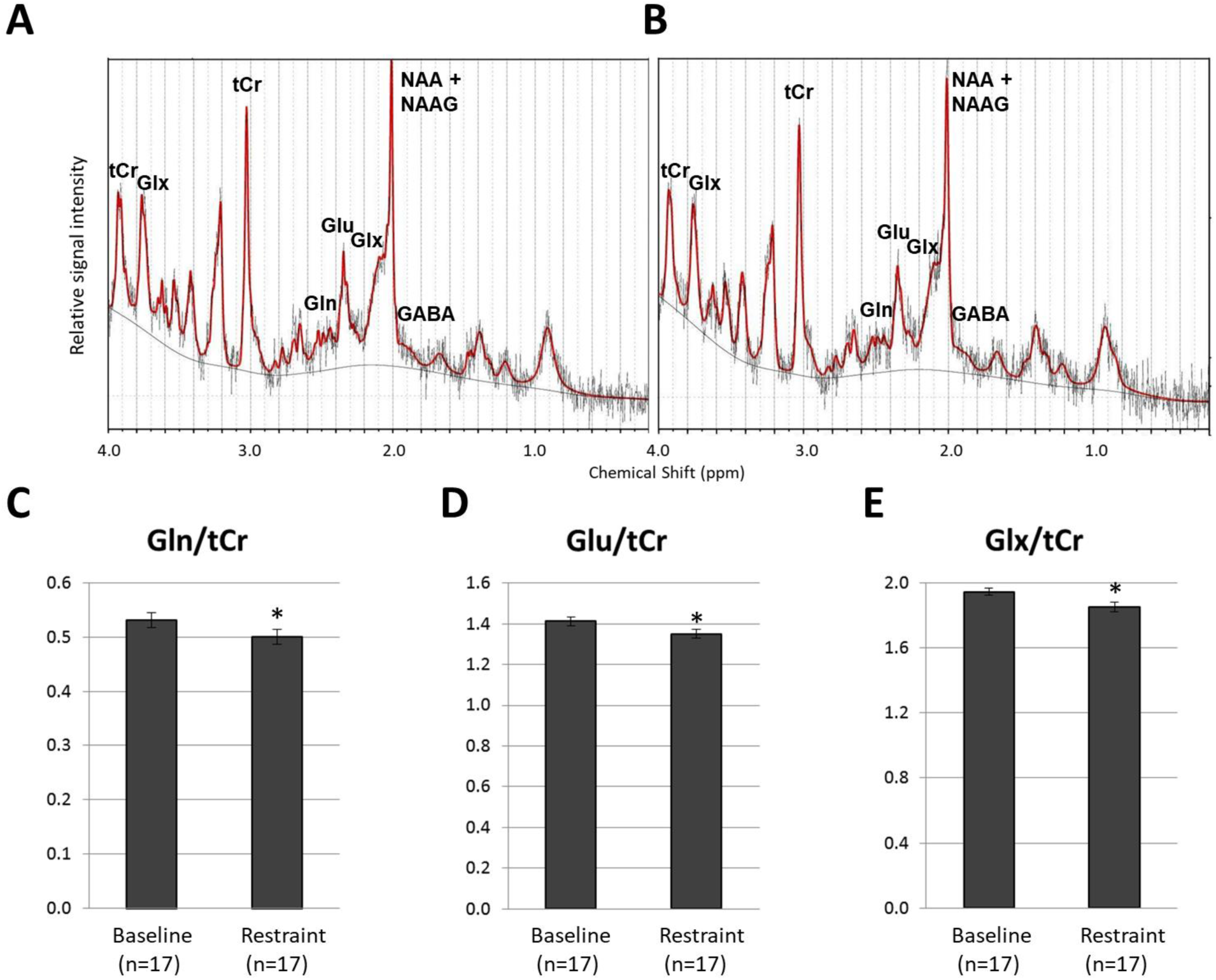
Representative spectra obtained from LCModel for proton magnetic resonance spectroscopy data for the baseline (A) and restraint groups (B) and effect of chronic restraint stress on glutamine (C), glutamate (D) and combined glutamine and glutamate (E). The figure shows spectra from a representative animal at baseline (A) and after 13 days of chronic restraint stress (B) depicting longitudinally reproducible peaks of various metabolites quantified using the LCModel. C-E show mean ± SEM of metabolite levels as a ratio to total creatine (tCr). Comparisons were made by a paired Wilcoxon signed-rank test (baseline, n=17; restraint, n=17). \**p* < 0.05.

### Change in hippocampal volume

Spearman’s rank correlation method revealed a significant correlation of whole-brain volumes with the weight of the animals at baseline (Fig. 6A), with the mean whole-brain volume to body weight ratio of the Sprague Dawley rats being 6.86 ± 0.16 mm^3^/g. A significant interaction effect was found between timepoint and group (Fig. 6B) and between timepoint, group and whole-brain volume (Table 2). This shows that the effect of timepoint on hippocampal volume was dependent on the group and whole-brain volume. Therefore, hippocampal volumes were normalised to whole-brain volumes for further analyses. A Pearson’s correlation test revealed no significant correlation between baseline percentage hippocampal volume and post-restraint latency to first immobility behaviour during FST (Fig. 6C). Percentage hippocampal volume decreased significantly following CRS (Fig. 6D) but did not change significantly in healthy controls (Table 2).

**Figure 6.**
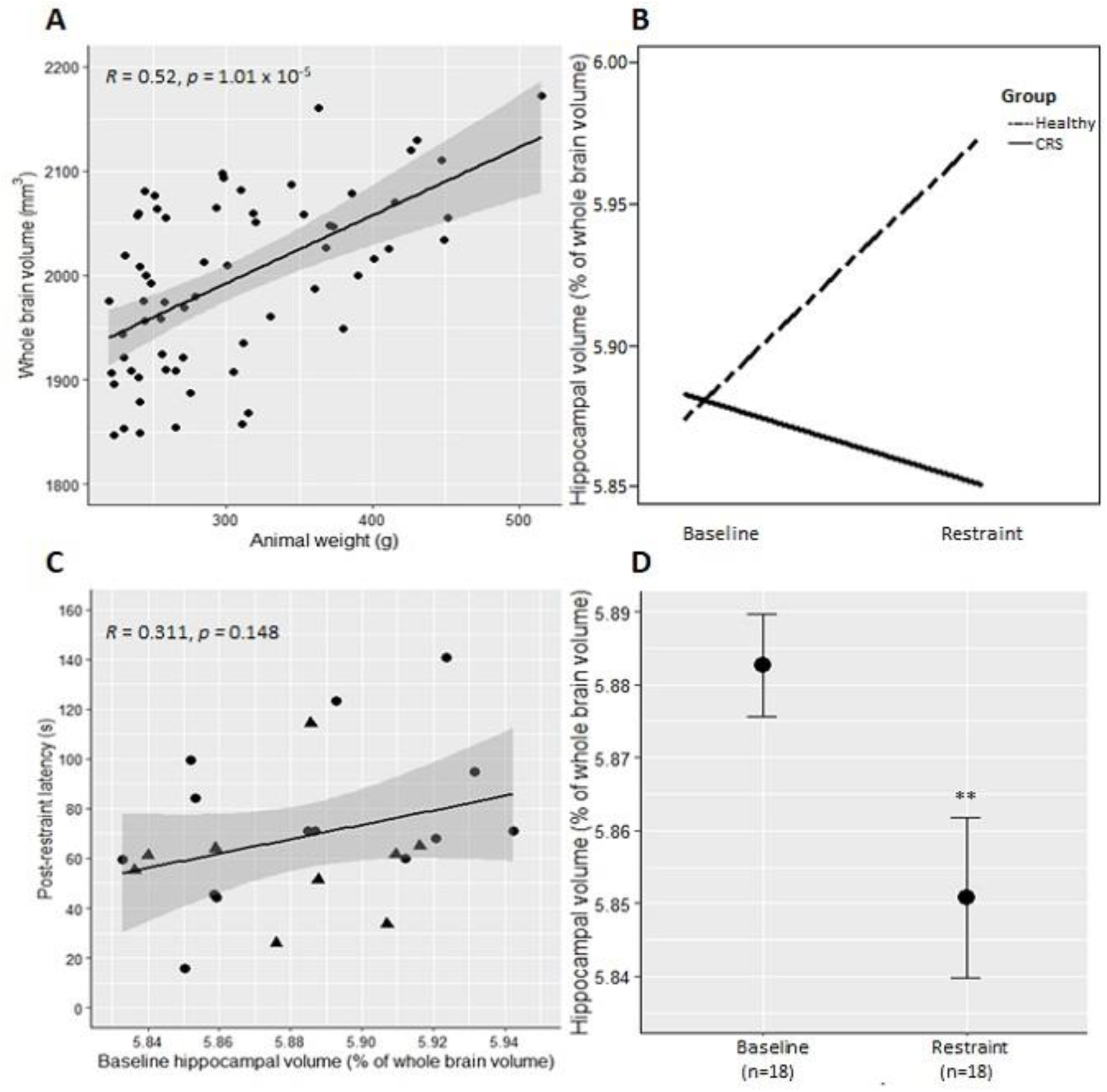
Correlation between weight of animals and whole-brain volume at baseline (A), interaction of timepoint (baseline vs restraint) and group (healthy vs with restraint) for percentage hippocampal volume (B), correlation between baseline percentage hippocampal volume and post-restraint latency to first immobility behaviour during forced swim test (C), and percentage hippocampal volume before and after chronic restraint stress (D) as measured from T2-weighted anatomical MRI data. A shows whole-brain volumes (mm^3^) plotted against the animal’s weight at baseline (n=65). Correlation was determined using Spearman’s rank correlation method. In B-D, hippocampal volumes were calculated as a percentage of whole-brain volume. B shows interaction between timepoint and group (healthy, n=8; restraint, n=18). The data represent mean of each group at each timepoint. C shows latency to first immobility behaviour during forced swim test plotted against hippocampal volume of the animals at baseline (baseline, n=23; restraint, n=23). Data points with triangular shape represent the nine animals which were used for FST-based ICA/dual regression analysis. D shows decrease in percentage hippocampal volume following 13 days of chronic restraint stress (baseline, n=18; restraint, n=18). The data represent mean ± SEM. Comparisons were made by paired t-test. **p < 0.01

### Correlations

Significant correlations were found between several behavioural and MRI measures (Fig. 7). However, none of the correlations survived multiple comparison correction, except for Glu/tCr with Glx/tCr (*p* < 0.0001) as would be expected.

**Figure 7.**
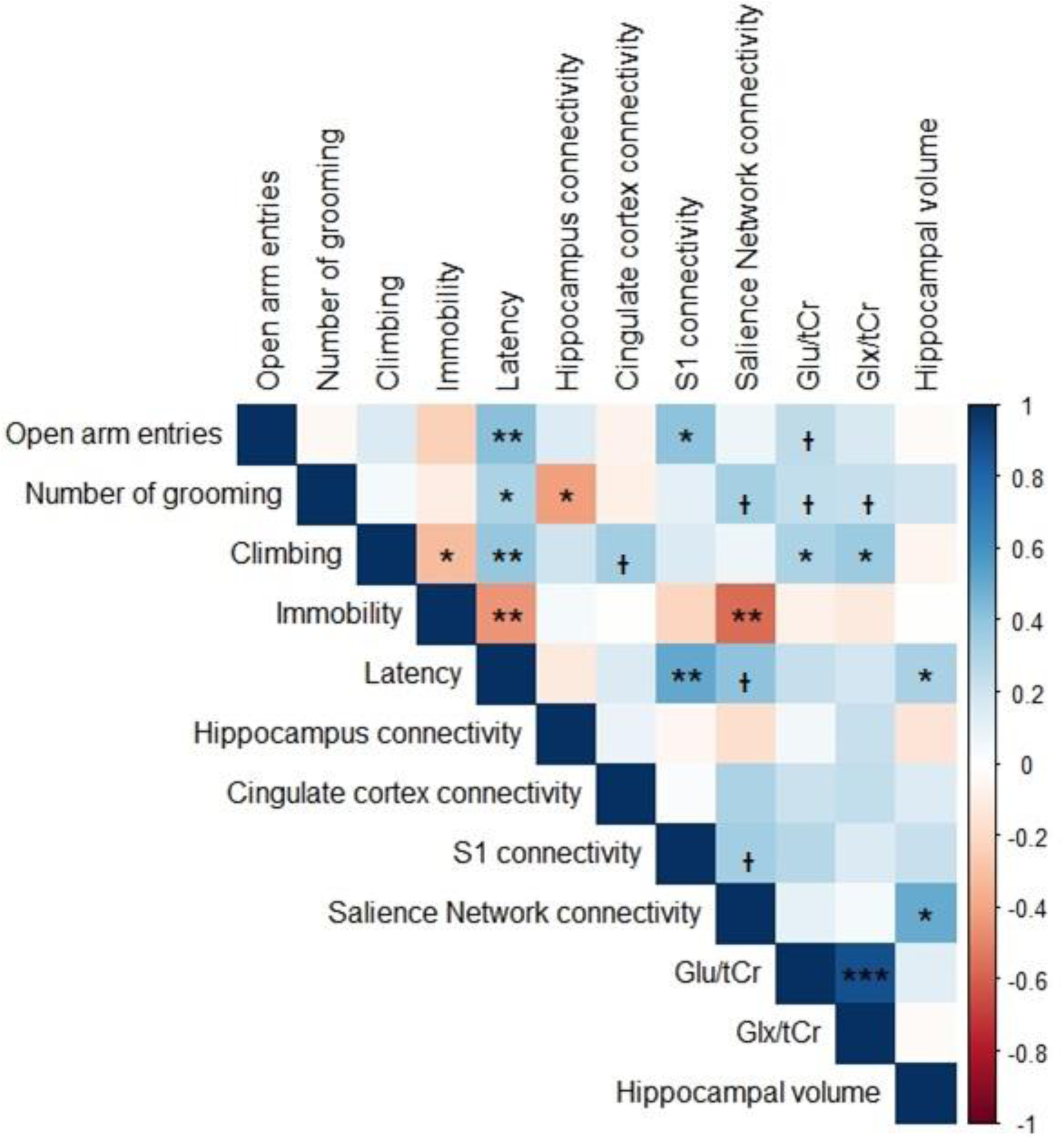
Correlations between behavioural tests and MRI measures. Comparisons of the following parameters were made by Pearson or Spearman correlations (depending on normality of data): open arm entries and number of grooming behaviours from elevated plus maze data; climbing score, immobility score and latency time from forced swim test data; connectivity (average z-scores) of the hippocampus, cingulate cortex, primary somatosensory cortex and salience network from the rs-fMRI data; Glu/tCr and Glx/tCr ratios from ^1^H-MRS data; and hippocampal volume. ᵻ *p* < 0.1, * *p* < 0.05, ** *p* < 0.01, *** *p* < 0.001 (no multiple comparison correction).

## Discussion

Animal models are an indispensable tool for studying aetiology, progress and treatment of depression in a controlled environment. However, there is still controversy regarding the validity of using rodent models of human neuropsychiatric disorders. Prior work in rodents investigating anxiety and depression-like behaviours (Suvrathan *et al.*, 2010; Ulloa *et al.*, 2010; Chiba *et al.*, 2012; Bogdanova *et al.*, 2013), peripheral biomarkers, functional connectivity of the brain (Henckens *et al.*, 2015) and hippocampal volume (Lee *et al.*, 2009; Alemu *et al.*, 2019) supports the validity of the CRS paradigm as a depression model (Wang *et al.*, 2017). However, our study is the first to correlate MRI measures of functional, chemical and structural changes in the brain with abnormal behaviour in the CRS model. Furthermore, similarities between our data and the MRI outcomes in humans suggest that the CRS model may be a useful component of translational studies aimed at developing and refining novel treatments for depression in humans.

### Aberrant resting-state functional connectivity following chronic restraint stress

ICA and dual regression analysis of our rodent rs-fMRI data detected decreased functional connectivity of the bilateral somatosensory cortex to the salience network, and of the right somatosensory cortex to the interoceptive network following CRS, and these reductions in connectivity were correlated with behavioural changes during FST and EPM. Altered functional connectivities in the equivalent human salience and interoceptive networks, which have an important role in being aware of, and orienting and responding to, biologically relevant stimuli have been identified in humans with depression and associated with their negative response biases (Harshaw, 2015). For example, in patients with bipolar disorder in the period of depression, Yin *et al.* (2018) observed decreased functional connectivity of insular cortex to somatosensory and motor cortex. In the present study, when a subset of animals showing the greatest depression-like behaviours in the FST was used in dual regression analysis, a decrease in connectivity of the right motor cortex and bilateral insular and somatosensory cortices to the salience network was also detected. In human studies, decreased functional connectivity in the insular cortex within the salience network in patients with depression is significantly correlated with the severity of symptoms (Manoliu *et al.*, 2014). Similarly, salience network connectivity was correlated with immobility behaviour during FST in our rats. CRS may induce abnormal behavioural responses in animals as a result of insular dysfunction within the salience network leading to an abnormal switching between the DMN and the central executive network (Manoliu *et al.*, 2014).

The DMN plays an important role in the pathophysiology of depression (Sheline *et al.*, 2009; Zhu *et al.*, 2012), being implicated in rumination, self-referential functions, and episodic memory retrieval (Lu *et al.*, 2012). One critical element of the DMN is the cingulate cortex, which has increased connectivity with other limbic areas in patients with depression (Greicius *et al.*, 2007; Sheline *et al.*, 2009; Davey *et al.*, 2012; Fang *et al.*, 2012; Rolls *et al.*, 2018). The results of the current study are consistent with these data: we detected hyperconnectivity of the cingulate cortex to the right retrosplenial cortex, visual cortex, inferior colliculus, bilateral thalamus, superior colliculus and hippocampus following CRS. However, cingulate cortex connectivity was not significantly correlated with behaviour or other MRI measures in our rats. This is surprising because cingulate cortex connectivity seems to play a role in clinical symptoms (Walter *et al.*, 2009), with higher functional connectivity leading to dysfunctional emotion, internal inspection, and endocrine regulation (Fang *et al.*, 2012). For example, increased functional connectivity between the thalamus and the cingulate cortex may result from increased emotional processing, at the cost of executive functions (Greicius *et al.*, 2007). However, our behavioural tests did not specifically address executive functioning in rats, and this could be addressed in future studies using appropriate cognitive tests.

Our rs-fMRI findings differ from a previous animal study using a shorter CRS protocol (2 h/day for 10 days), which did not find any significant changes in the RSNs despite using the same ICA/dual regression approach of rs-fMRI data analysis performed here (Henckens *et al.*, 2015). Moreover, when comparing ‘overall connectivity strength’, connectivity was increased in somatosensory and visual networks, which was not observed in our experiments. The shorter duration of the restraint stress, as well as intrinsic differences between the rs-fMRI data analysis methods used to detect changes in connectivity, and the effect of an isoflurane-only anaesthetic protocol on RSNs (Paasonen *et al.*, 2018) used in the previous study (Henckens *et* al., 2015) could be the cause of these inconsistencies.

It is interesting to compare the hyperconnectivity observed here to findings in other animal models used to investigate depression and anxiety. Brain activation in cortical and hippocampal regions in mice following chronic social defeat stress is observed in manganese-enhanced magnetic resonance imaging (Laine *et al.*, 2017). Additionally, aberrant hippocampal, thalamic and cortical connectivity is reported in other rodent models using different data acquisition and/or analysis methods. For example, in the chronic unpredictable stress rat model, rs-fMRI studies found increased functional connectivity of the hippocampus to several brain regions (Magalhães *et al.*, 2019), increased functional connectivity between atrophied brain regions such as the hippocampus, striatum and cingulate, motor and somatosensory cortices (Magalhães *et al.*, 2018) and increased regional homogeneity (coherence of intraregional spontaneous low-frequency activity) in the hippocampus, thalamus and visual cortex as well as a decreased regional homogeneity in the motor cortex (Li *et al.*, 2018). Electrophysiology studies have also reported long-lasting inhibition of long-term potentiation in the thalamo-cortical circuitry (Zheng *et al.*, 2012) and in the hippocampal–cortical circuitry (Cerqueira *et al.*, 2007) in chronic unpredictable stress models while hippocampal–cortical circuitry inhibition was also reported in acute platform stress rat models (Rocher *et al.*, 2004). These different animal models reflect specific aspects of depression and therefore, they may be useful for understanding the heterogeneity of human depression.

### Decrease in glutamate and glutamine levels following chronic restraint stress

Another major finding of this study was the significant decreases in Gln, Glu and Glx in the left sensorimotor cortex following CRS. Both animal and human studies suggest that glutamatergic neurotransmission plays a pivotal role in the depression pathophysiology (Marrocco *et al.*, 2014; Moriguchi *et al.*, 2019). Human ^1^H-MRS studies have reproducibly reported a reduced concentration in Glu, Gln and/or Glx in several brain regions including the anterior cingulate cortex (Mirza *et al.*, 2004; Luykx *et al.*, 2012) and the prefrontal cortex (Hasler *et al.*, 2007; Portella *et al.*, 2011). Similarly, other ^1^H-MRS studies of animal models of depression such as the chronic mild stress and the chronic social isolation models have reported decreases in these neurometabolites in the prefrontal cortex (Hemanth Kumar *et al.*, 2012) and hippocampus (Hemanth Kumar *et al.*, 2012; Shao *et al.*, 2015). The correlation between decreasing Glu and Glx levels and reduced climbing behaviour during FST suggests that the shortage in these neurometabolites might be due to a reduction in the number of astrocytes which in turn alters neuronal activity and therefore may contribute to depression-like behaviours, as previously shown in an L-α aminoadipic acid (L-AAA) infusion mouse model (Lee *et al.*, 2013).

### Decrease in hippocampal volume

A reduction in hippocampal volume has been consistently associated with depression in humans (McKinnon *et al.*, 2009). However, the stage at which hippocampal atrophy begins in human depression is unclear and so is the direction of causality. There are two main hypotheses regarding how depression is associated with hippocampal atrophy. Firstly, hippocampal volume reduction, probably as a result of early life adversity, poverty and stress, might predispose people to depression. This hypothesis seems consistent with smaller hippocampal volumes already present in first depressive episodes (Cole *et al.*, 2011) and in young children (Barch *et al.*, 2019) and adolescents (Rao *et al.*, 2010) with depression. The second hypothesis, known as the neurotoxicity hypothesis, suggests that cumulative exposure to disrupted emotion regulation, stress reactivity, glucocorticoids and antidepressant medications as a result of depression increases neuronal susceptibility to insults and therefore leads to hippocampal deficits (Sheline, 2011). This hypothesis is consistent with hippocampal atrophy being more pronounced among individuals with recurrent episodes and in chronic depression (McKinnon *et al.*, 2009; Cheng *et al.*, 2010; Brown *et al.*, 2014). The longitudinal nature of the present study precludes the first hypothesis in CRS animals. Hippocampal volume decreased after CRS, while animals which did not undergo CRS did not have any significant change in hippocampal volume. This study supports the neurotoxicity hypothesis and further suggests that the reduction in hippocampal volume might happen at a very early stage in depression, within only three weeks in this animal model.

### Study limitations

Our study has four main limitations. Firstly, only male rats were used in this study, even though CRS has been shown to successfully induce depression-like behaviours in freely cycling adolescent female rats (Hibicke *et al.*, 2017a; Hibicke *et al.*, 2017b). Future studies could expand the applicability of present results by investigating brain changes following CRS in female rats. Secondly, the SPT did not detect anhedonia in our animals following restraint, despite anhedonia being a well-documented effect of CRS (e.g., Chiba *et al.*, 2012; Ampuero *et al.*, 2015; Liu *et al.*, 2016). Use of non-acidified water and longer habituation and/or test times as performed in these studies may be required. Thirdly, MRI data was acquired under anaesthesia, which could potentially alter the blood-oxygen-level-dependent (BOLD) signal detection. However, functional connectivity patterns of animals anaesthetised using a combination of low-dose isoflurane and medetomidine have good correspondence with those of awake rats (Paasonen *et al.*, 2018) with strong inter-cortical and cortical-subcortical functional connectivity (Grandjean *et al.*, 2014; Bukhari *et al.*, 2017) and are reproducible (Lu *et al*., 2012). Moreover, ^1^H-MRS data was acquired only in the left sensorimotor cortex. Future studies can investigate neurometabolite changes in bilateral sensorimotor cortex as well as in other brain regions such as the basal ganglia, hippocampus, anterior cingulate cortex, and occipital cortex, which are extensively investigated in ^1^H-MRS studies of human depression. Neurometabolite and structural changes could be confirmed using invasive methods following CRS. Finally, the pharmacological or interventional validity of the present neuroimaging findings is unknown. Future work should examine the utility of these findings as preclinical target engagement biomarkers with pharmacological and neuromodulatory interventions. If this proves to be the case, this animal model has potential utility for high throughput dose finding studies of neurotherapeutics and novel interventions.

### Conclusion

The present study is the first to demonstrate significant changes in functional connectivity, neurometabolite levels and hippocampal volume in the same animals post-CRS and the correlation of these measures with changes in behaviour provide insight into the neurobiological changes that may underpin patient symptoms. Cumulative exposure to stress might increases neuronal and astrocytic death leading to hippocampal atrophy and a shortage in glutamate and glutamine, which in turn alters neuronal activity and therefore contribute to learned helplessness. The lack of correlation of depression-like behaviours with cingulate cortex connectivity was surprising but increased cingulate cortex connectivity may instead be negatively correlated with executive functioning, which was not tested in the present study. Overall, the substantial concordance of the present findings with the human literature of depression presents a unique opportunity for the integration of behavioural, cellular and molecular changes detected in this animal model of depression when testing the effects of new drug treatments or therapies with changes in MRI measures of brain function, chemistry and structure that may be translated to future studies of the human disorder.

## Funding and Disclosure

This research was funded by The University of Western Australia. BJS is supported by a Forrest Research Foundation Scholarship, an International Postgraduate Research Scholarship, and a University Postgraduate Award. LAH is supported by the Commonwealth Government’s ‘Australian Government Research Training Program Fees Offset’. KWF was an Australian National Imaging Facility Fellow, a facility funded by the University, State and Commonwealth Governments. PEC was supported by the National Institute of Mental Health Grant R01 MH113700. JR was supported by a Fellowship from MSWA and the Perron Institute for Neurological and Translational Science.

The authors declare no potential conflicts of interest with respect to the research, authorship, and/or publication of this article.

## Acknowledgements

The authors thank Ms Marissa Penrose-Menz, Ms Kerry Leggett, Ms Elizabeth Jaeschke-Angi, Ms Leah Mackie, Ms Kaylene Schutz, Ms Yashvi Bhatt and Mr Parth Patel for their assistance with the experiments and the team at M-Block Animal Care Services, especially Ms Sandra Goodin and Mr Stefan Davis, for their assistance with animal care and provision of some essential materials for the behavioural experiments. The authors acknowledge the facilities and scientific and technical assistance of the National Imaging Facility, a National Collaborative Research Infrastructure Strategy (NCRIS) capability, at the Centre for Microscopy Characterisation and Analysis, The University of Western Australia. The content is solely the responsibility of the authors and does not necessarily represent the official views of the National Institutes of Health.

## Author contributions

BJS and JR designed the research. BJS performed the experiments and analysed the SPT, FST, rs-fMRI, ^1^H-MRS and structural MRI data. LAH helped with the experiments and analysed the EPM data. KWF provided troubleshooting and methodological advice on acquiring and analysing the MRI data. PEC provided troubleshooting and methodological advice on the analysis of rs-fMRI and structural MRI data. BJS wrote the first draft of the paper. BJS, KWF, SJE, PEC and JR assisted with the editing and revision of subsequent drafts of the manuscript.

